# Structural and Dynamic Basis of TREM2–DAP12 Stabilization by Small-Molecule Agonist VG-3927

**DOI:** 10.64898/2026.06.22.732806

**Authors:** Fanyu Zhao, Wei Xia, Xintai Zhang, Xue Wu, Junxiao Ding, Xiangyue Li, John Z. H. Zhang

**Affiliations:** NYU-ECNU Center for Computational Chemistry and Shanghai Frontiers Science Center of Artificial Intelligence and Deep Learning, NYU Shanghai, Shanghai 200124, China; Shanghai Engineering Research Center of Molecular Therapeutics and New Drug Development, Shanghai Key Laboratory of Green Chemistry & Chemical Process, School of Chemistry and Molecular Engineering, East China Normal University at Shanghai, 200062, China; Faculty of Synthetic Biology, Shenzhen University of Advanced Technology, Shenzhen 518107, China; Department of Chemistry, New York University, New York, New York 10003, U.S.A; Key Laboratory of Quantitative Synthetic Biology, Shenzhen Institute of Synthetic Biology, Shenzhen Institutes of Advanced Technology, Chinese Academy of Sciences, Shenzhen 518055, China; Collaborative Innovation Center of Extreme Optics, Shanxi University, Taiyuan 030006, China

**Keywords:** Alzheimer’s disease, TREM2, DAP12 dimer, VG-3927, transmembrane binding, membrane-associated binding, ASGBIE

## Abstract

Triggering receptor expressed on myeloid cells 2 (TREM2) is an important microglial receptor implicated in Alzheimer’s disease, but the structural basis of small-molecule TREM2 agonism remains poorly understood. In particular, the clinical-stage agonist VG-3927 has been reported to promote TREM2–DAP12 complex formation, suggesting a mechanism distinct from conventional recognition at the ectodomain surface. Here, we propose that VG-3927 binds within a membrane-embedded interfacial cavity formed by the transmembrane helices of TREM2, DAP12A, and DAP12B. Our simulations support a stable and reproducible cavity-bound pose that is maintained across independent trajectories and is anchored by a persistent hotspot network dominated by hydrophobic and van der Waals interactions. Rather than merely occupying a pre-existing pocket, VG-3927 reshapes the surrounding transmembrane assembly by reinforcing selected interhelical contacts, strengthening receptor–adaptor hydrogen-bond coupling, and biasing the three-helix bundle toward a more compact, more upright, and more conformationally focused state. Importantly, the ligand-bound ensemble remains within a subdomain of the apo conformational landscape, consistent with stabilization of a pre-existing but less populated signaling-relevant state. Together, these findings support a membrane-asscosiated mechanism in which VG-3927 acts as a transmembrane interfacial *molecular glue*, biasing the TREM2–DAP12 complex toward a more compact, ordered, and signaling-competent state. This finding can provide a structural framework for understanding small-molecule TREM2 agonism.

## 1 INTRODUCTION

Triggering receptor expressed on myeloid cells 2 (TREM2) is a type I transmembrane immune receptor of the immunoglobulin superfamily. In the central nervous system, TREM2 is primarily expressed by microglia and functions as an important regulator of innate immune sensing.^1–4^ In Alzheimer’s disease (AD), TREM2 has emerged as a key modulator of disease-associated microglial states, influencing microglial activation and survival, lipid sensing, amyloid-β (Aβ) plaque responses, phagocytosis, and tau-associated pathology.^1,2,5^ Human genetic studies have further showed that rare TREM2 variants with reduced function, particularly R47H and R62H, increase the risk of late AD, establishing TREM2 as a biologically relevant risk gene and a potentially target in AD treatment.^3,4,6^

Mature TREM2, after removal of the N-terminal signal peptide, consists of an extracellular domain (ECD), a single-pass transmembrane (TM) helix, and a short cytoplasmic tail.^7,8^ The ECD contains a V-type immunoglobulin-like (Ig-like) domain involved in ligand recognition and a flexible stalk region that undergoes proteolytic shedding to generate soluble TREM2 (sTREM2). Structural studies indicate that TREM2 does not contain a classical deep ligand-binding pocket. Instead, ligand recognition is mediated by several shallow surfaces in the Ig-like domain. One major site is a hydrophobic surface near the complementarity-determining region-like (CDR-like) loops, which binds to disease-relevant ligands such as apolipoprotein E (ApoE) and Aβ-related species.^7,9,10^ A second site is a positively charged basic surface that includes residues such as R47 and R62 and participates in the recognition of anionic lipids, glycans, and other negatively charged ligands.^7,9,10^ These extracellular recognition events, however, cannot be fully understood without considering the membrane-embedded receptor–adaptor assembly that link ligand engagement with intracellular signaling.

TREM2 has a short cytoplasmic tail and lacks intrinsic catalytic or canonical signaling motifs. Signal transduction therefore depends on its association with DNAX-activating protein of 12 kDa (DAP12, also known as TYROBP), a disulfide-linked adaptor dimer containing immunoreceptor tyrosine-based activation motifs (ITAMs) in its cytoplasmic domains.^1,11^ The TREM2–DAP12 complex is stabilized by salt-bridge like interactions between a conserved lysine residue in the TREM2 TM helix and aspartate residues in the DAP12 TM helices.^11–13^ Through this membrane-associated coupling, extracellular ligand recognition by TREM2 promotes DAP12 phosphorylation and downstream recruitment of SYK and related signaling pathways. Thus, the TM region is not simply a membrane anchor but an essential structural element that couples extracellular sensing to intracellular immune signaling.

Because of its central role in microglial biology, TREM2 has become an active target for therapeutic development. Most clinical strategies have focused on agonistic antibodies that bind the TREM2 ECD.^14^ AL002, one of the most advanced TREM2 agonistic antibodies, showed central target engagement but did not meet the primary clinical efficacy endpoint in a Phase 2 study in early AD.^15^ DNL919 was discontinued during clinical development based on emerging early-phase data,^16^ whereas VHB937 remains under clinical evaluation, including a Phase 2 study in early AD.^17^ These outcomes highlight both the therapeutic relevance and the biological complexity of TREM2 modulation.

In parallel with antibody-based approaches, small-molecule TREM2 agonists have attracted increasing attention because they may offer oral administration, improved brain exposure, and reduced dependence on high peripheral antibody concentrations. VG-3927, originally developed by Vigil Neuroscience and subsequently acquired by Sanofi, is a clinical-stage, orally bioavailable, brain-penetrant small-molecule TREM2 agonist.^18,19^ Publicly disclosed preclinical and Phase 1 data describe VG-3927 as a potent and selective TREM2 agonist with positive allosteric activity. It enhances TREM2 responses in cooperation with endogenous damage-associated ligands and is insensitive to the presence of sTREM2, suggesting a mechanism distinct from direct engagement of the soluble ECD.^18–20^ Additional disclosed data indicate that VG-3927 promotes DAP12 and SYK phosphorylation and increases TREM2–DAP12 complex formation in cellular assays, supporting a molecular-glue-like mechanism that stabilizes productive receptor–adaptor assemblies.^21^

Despite these advances, the structural basis by which VG-3927 enhances TREM2 signaling remains unresolved. The absence of detectable binding to sTREM2 suggests that the small molecule may require the membrane-bound receptor context. Given the dependence of TREM2 signaling on TM helix association with DAP12, the TM interface represents a plausible and mechanistically relevant site for small-molecule modulation. In particular, a lipophilic molecule such as VG-3927 could, in principle, insert into the membrane and stabilize interhelical cavities formed within the TREM2–DAP12 assembly. This hypothesis is consistent with the reported enhancement of receptor–adaptor complex formation, although direct experimental structural evidence for a TM binding site is not yet available.

Here, the objective of the present study was to characterize the potential membrane-associated binding mode of VG-3927 and to evaluate its structural effects on the TREM2–DAP12 transmembrane assembly. Because no experimental structure of the ligand-bound full-length complex is currently available, we constructed a structural model of the TREM2–DAP12 complex using AlphaFold 3 and embedded the assembly in an explicit phospholipid bilayer. We then performed blind docking with AutoDock Vina to explore possible binding regions in the transmembrane environment, followed by all-atom molecular dynamics (MD) simulations to refine and assess protein–ligand interactions. Our simulations indicate that VG-3927 can stably occupy a wedge-shaped interfacial cavity formed by the TM helices of TREM2 and the DAP12 dimer. Free-energy decomposition using alanine scanning with generalized Born and interaction entropy (ASGBIE) identified key residues that contribute to ligand stabilization within this membrane-embedded site. Furthermore, analyses of interhelical contact occupancies and helix-packing geometry suggest that VG-3927 binding promotes a more compact and structurally reinforced TREM2–DAP12 interface.

Together, these findings provide a structural model for membrane-associated small-molecule modulation of the TREM2–DAP12 complex. By suggesting how a lipophilic agonist may stabilize an interhelical receptor–adaptor interface, this study offers a mechanistic framework for understanding VG-3927 action and for guiding the structure-based optimization of next-generation TREM2 modulators. More broadly, our results support the concept that transmembrane interfaces in immune receptor complexes can serve as pharmacologically accessible sites for small-molecule regulation.

## 2 RESULT AND DISCUSSION

### 2.1 Structure of TREM2-DAP12 complex

**Figure 1.**
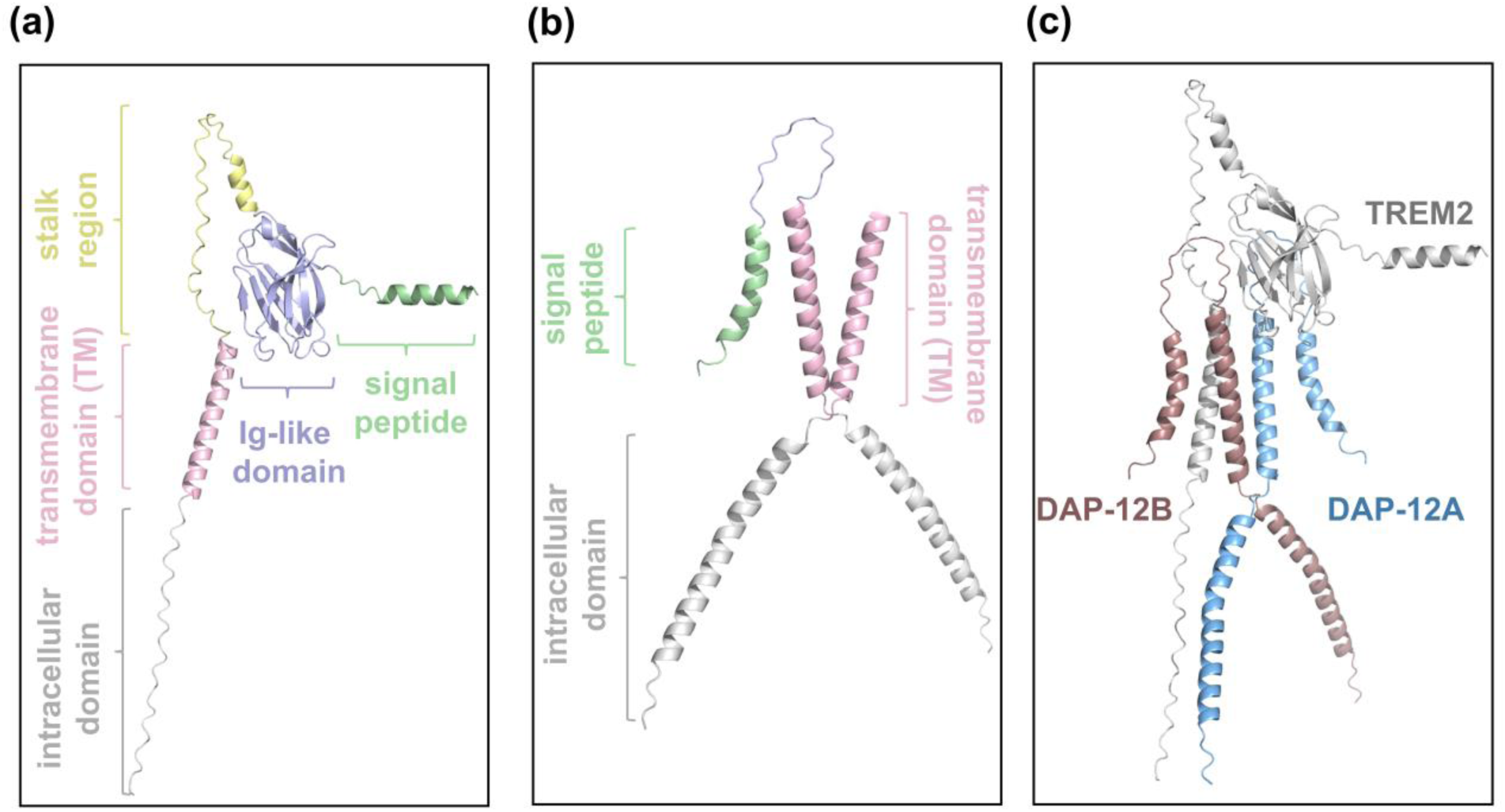
Structural overview of the full-length TREM2 (a), DAP12 dimer (b) and TREM2-DAP12 complex (c), with functional structures colored differently.

To obtain a structurally reasonable starting model for the membrane-associated TREM2–DAP12 complex, we used AlphaFold 3 to predict the receptor–adaptor assembly and then generated a trimmed construct for MD simulations (**Figure 2**). The signal peptides and cytoplasmic regions of TREM2 and DAP12 were removed to reduce computational cost and to focus the simulations on the extracellular juxtamembrane and TM regions. For DAP12, the extracellular loop was further truncated while retaining the minimal N-terminal region required to preserve the two interchain disulfide bonds. This strategy reduced the influence of highly flexible regions distant from the transmembrane interface while maintaining the structural elements required for a realistic TREM2–DAP12 receptor–adaptor architecture.

**Figure 2.**
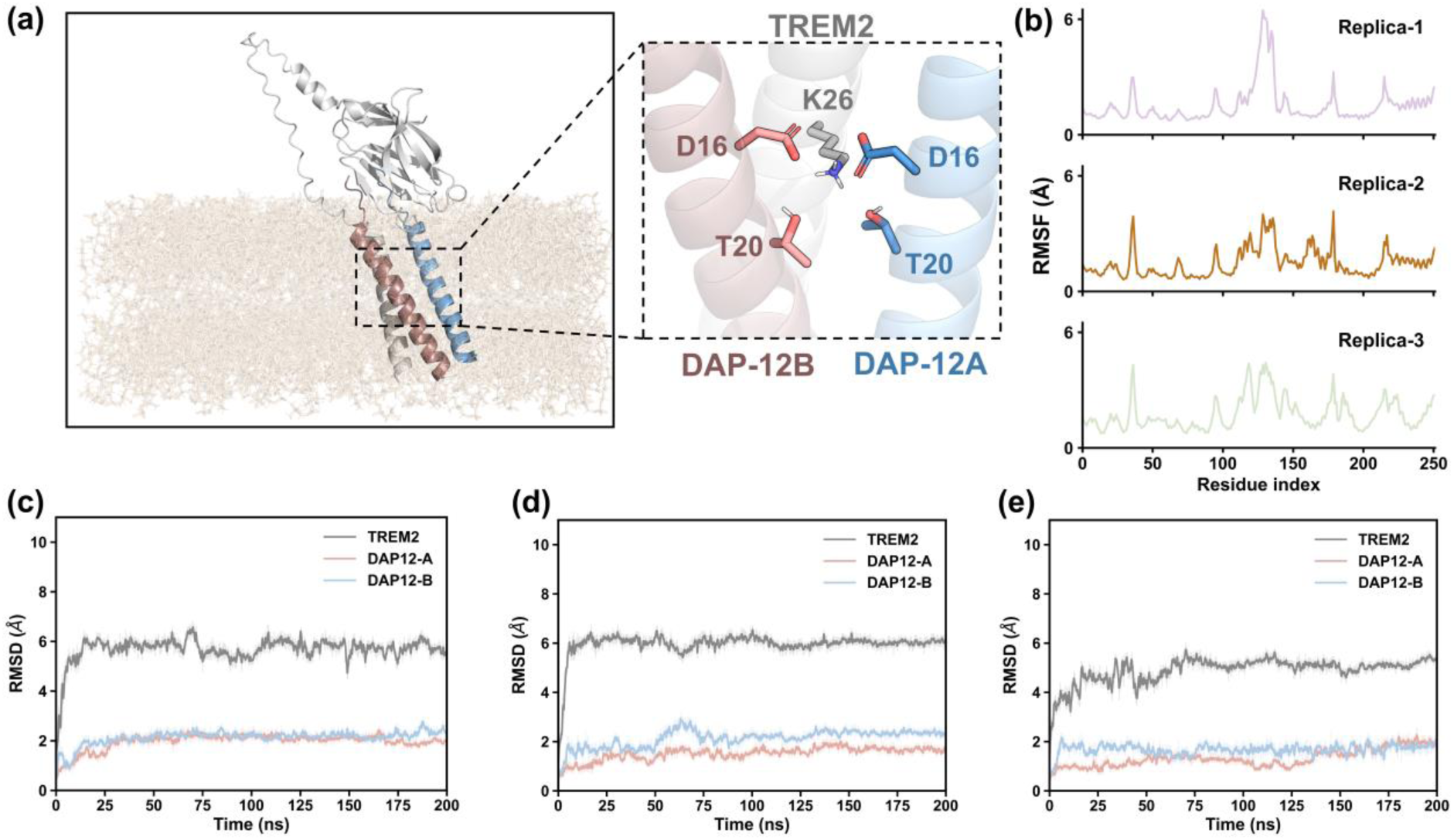
(a) Predicted structure of the TREM2–DAP12 complex embedded in a membrane, with an enlarged view of the conserved transmembrane anchoring network shown without the membrane for clarity. TREM2, DAP12A, and DAP12B are colored gray, blue, and red, respectively, and the membrane is shown as yellow sticks. Residues involved in the conserved anchoring network are labeled according to the literature numbering, whereas subsequent trajectory analyses use the continuous simulation-specific numbering described in the text. (b) RMSF profiles from three independent replicas. (c–e) RMSD profiles of TREM2, DAP12A, and DAP12B in replica 1, replica 2, and replica 3, respectively, during 200 ns MD simulations. TREM2, DAP12A, and DAP12B are shown in gray, red, and blue, respectively.

The resulting model captured the characteristic three-helix organization of the TREM2–DAP12 TM complex, with the TREM2 helix positioned between the two DAP12 helices (**Figure 2a**). Importantly, the predicted interface contained the conserved polar network previously reported for this receptor–adaptor complex. In this paragraph, residues in the anchoring network are described using the literature numbering to facilitate comparison with previous studies. The network involves TREM2 K26 and DAP12 D16/T20 residues from both adaptor helices. These interactions include salt-bridge-like electrostatic contacts between TREM2 K26 and DAP12 D16 residues, together with additional hydrogen-bonding contacts involving DAP12 T20. The presence of this interhelical interaction network supports the structural plausibility of the AlphaFold 3 model. The overall three-helix arrangement is also consistent with recent membrane-associated structural studies of the TREM2–DAP12 transmembrane complex, which support a receptor–adaptor assembly stabilized by conserved polar interactions within the membrane.^12^

Subsequent MD simulations further supported the stability of the starting model. Across three independent 200 ns replicas, the root-mean-square fluctuation (RMSF) profiles showed similar residue-dependent patterns, indicating reproducible conformational dynamics (**Figure 2b**). The root-mean-square deviation (RMSD) trajectories further showed that both DAP12 helices remained stable throughout the simulations, whereas TREM2 exhibited moderately higher RMSD values (**Figure 2c–e**). This increase was mainly attributable to the flexible extracellular stalk region rather than disruption of the transmembrane core. Together, the conserved interhelical anchoring network, stable RMSD behavior of the DAP12 dimer, and reproducible RMSF profiles support the use of this AlphaFold 3-derived model for subsequent docking and mechanistic analyses.

For clarity in trajectory analysis, all residues were renumbered continuously in the simulation system. In this numbering scheme, TREM2 residues 1–179 correspond to residues 21–199 in the standard sequence, whereas DAP12A residues 180–215 and DAP12B residues 216–251 both correspond to DAP12 residues 3–38. Unless otherwise stated, all residue indices reported below refer to this simulation-specific numbering.

### 2.2 Predicted complex structure of VG-3927 binding to TREM2-DAP12

To investigate the stability and reproducibility of VG-3927 binding within the TM region of the TREM2–DAP12 complex, we employed a blind docking protocol followed by explicit membrane MD validation. A representative receptor conformation was extracted from the preceding 200 ns apo-state MD simulations and used for docking with AutoDock Vina. A high-scoring pose (-9.6 kcal/mol) located within an interhelical cavity was selected as the initial binding model for subsequent simulations. To evaluate the convergence and stability of this binding mode, three independent 200 ns replicas were initiated from the same docked pose in an 80:20 POPC:cholesterol bilayer. Additional details of the simulation setup are provided in the Methodology section.

As shown in **Figure 3a**, VG-3927 remained located within an interfacial cavity formed by the TM helices of TREM2, DAP12A, and DAP12B throughout the simulations. To assess whether the initial docking pose leads to a reproducible bound state, we compared four representative ligand conformations: the centroid structures obtained from k-means clustering of each individual replica trajectory (i.e., purple, yellow and green stick models in **Figure 3a**) and the centroid structure obtained from clustering the concatenated trajectory of all three replicas (i.e., silver stick models in **Figure 3a**). These representative structures showed strong spatial overlap within the TM cavity and adopted similar packing orientations against the surrounding helices. The convergence of ligand placement across independent simulations suggests that the selected docking pose represents a stable binding mode rather than a stochastic event.

**Figure 3.**
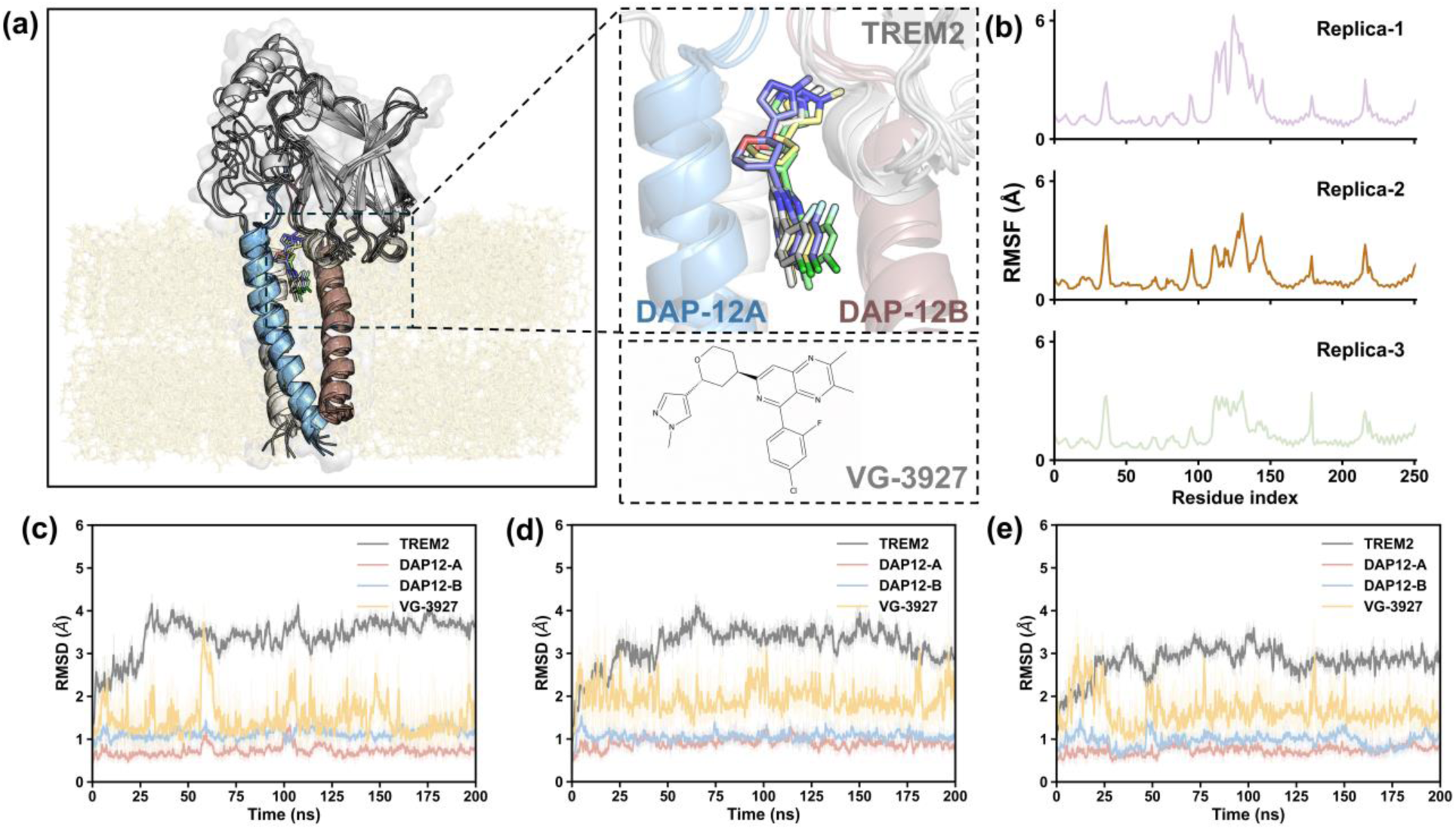
(a) Membrane-embedded structure of the VG-3927-bound TREM2–DAP12 complex, with an enlarged view of the transmembrane binding site shown without the membrane for clarity. Representative ligand poses from replicas 1, 2, and 3 are shown in purple, yellow, and green, respectively, whereas the representative pose obtained from the concatenated trajectories is shown in silver. The membrane is shown as yellow sticks, and the 2D structure of VG-3927 is shown below the enlarged binding-site view. (b) RMSF profiles from three independent replicas. (c–e) RMSD profiles of TREM2, DAP12A, DAP12B, and VG-3927 in replica 1, replica 2, and replica 3, respectively, during 200 ns MD simulations. TREM2, DAP12A, DAP12B, and VG-3927 are shown in gray, red, blue, and yellow, respectively.

The RMSD profiles further support the stability of the ligand-bound complex (**Figure 3c–e**). Across the three replicas, VG-3927 showed limited fluctuations and remained largely within the same binding cavity, with ligand RMSD values generally maintained in the low-angstrom range and only transient fluctuation. The protein scaffold also remained stable during the simulations. DAP12A and DAP12B displayed consistently low RMSD values, indicating stable TM helix organization, whereas TREM2 showed moderately higher RMSD values, mainly reflecting the greater mobility of its extracellular stalk region rather than disruption of the TM interface. Consistently, RMSF profiles revealed similar residue-dependent fluctuation patterns among the three replicas (**Figure 3b**). Although local variations were observed, the TM regions exhibited relatively low fluctuations, indicating that VG-3927 is accommodated within a structurally stable and dynamically reproducible membrane-embedded environment. The major high-fluctuation regions can be assigned to structurally flexible or solvent-exposed segments of the TREM2–DAP12 assembly. Residues 36–38 are located in the Ig-like domain, within the connecting β-strand between the CDR1 and CDR2-like loops. Residues 128–137 are close to the ADAM10/17-mediated shedding site in the flexible stalk region, where higher flexibility is expected. In addition, residues 178–179 and 215–216 correspond to the N-terminal extracellular loop regions of the two DAP12 helices in the trimmed simulation construct. These regions are located outside the tightly packed transmembrane core and are more exposed to solvent. Therefore, the increased RMSF values observed in these segments are structurally reasonable.

The physicochemical properties of VG-3927 are also consistent with localization in a membrane-associated cavity. SwissADME^22^ and ADMETlab^23^ predictions indicate low aqueous solubility and substantial lipophilicity. SwissADME predicted a consensus LogP_o/w_ of 4.29, with individual models ranging from 2.78 to 5.57, and LogS values as low as -8.60 using the SILICOS-IT model. ADMETlab similarly predicted a LogS of -5.12, LogP of 3.90, and LogD7.4 of 3.34. Together, these values characterize VG-3927 as a hydrophobic small molecule with a strong tendency for lipid or membrane-associated binding. Structurally, its fused aromatic heterocyclic scaffold and halogen substituents further support favorable accommodation within a lipid-exposed or protein–membrane interfacial environment rather than a highly solvent-exposed surface. These properties support the use of a membrane-embedded receptor model and justify exploration of the TM region by blind docking.

Overall, these results indicate that the VG-3927-bound pose is sufficiently stable for residue-level free-energy decomposition and contact-network analysis. Because the ligand occupies an interfacial cavity jointly formed by TREM2 and DAP12, it is positioned not only to establish direct protein–ligand contacts but also to influence TM helix packing at the receptor–adaptor interface. This provides the structural basis for the subsequent analysis of binding hotspots and ligand-induced remodeling of the TREM2–DAP12 TM assembly.

### 2.3 Hotspot analysis of VG-3927 binding to TREM2-DAP12

We next performed per-residue free-energy decomposition using ASGBIE to identify the residues that stabilize VG-3927 within the TM cavity. As shown in **Figure 4a**, the free-energy averaged results from three independent replicas identified nine major hotspot residues located within 5 Å of the VG-3927 binding site and contributing more than 1 kcal·mol^−1^ to ligand stabilization. These residues include F54, I155, and L159 from TREM2; I229 from DAP12B; and I193, V185, L190, D197, and V194 from DAP12A, which reveals an asymmetric but cooperative binding pattern across the three-helix interface. Although all three helices participate in ligand recognition, their roles are not equivalent. F54 and I155 from TREM2 and I229 from DAP12B showed the largest contributions, each exceeding 2 kcal·mol^−1^, indicating that they serve as the principal energetic anchors of the ligand-bound complex. The remaining residues in DAP12A forms an extended hydrophobic wall, contributing between 1 and 2 kcal·mol^−1^ and collectively forming a secondary stabilizing shell around the ligand. These results suggest that VG-3927 binding is supported by several energetically important residues rather than by diffuse weak contacts across the entire binding pocket. Also, this spatial distribution of hotspots residues indicates that VG-3927 does not simply bind to one protein surface; instead, it bridges the receptor and adaptor helices within the TM cavity. Such a binding mode is consistent with a molecular-glue-like mechanism in which the ligand stabilizes the TREM2–DAP12 assembly by reinforcing the interhelical interface. Detailed information is provided in **Table S1**-**S3**.

**Figure 4.**
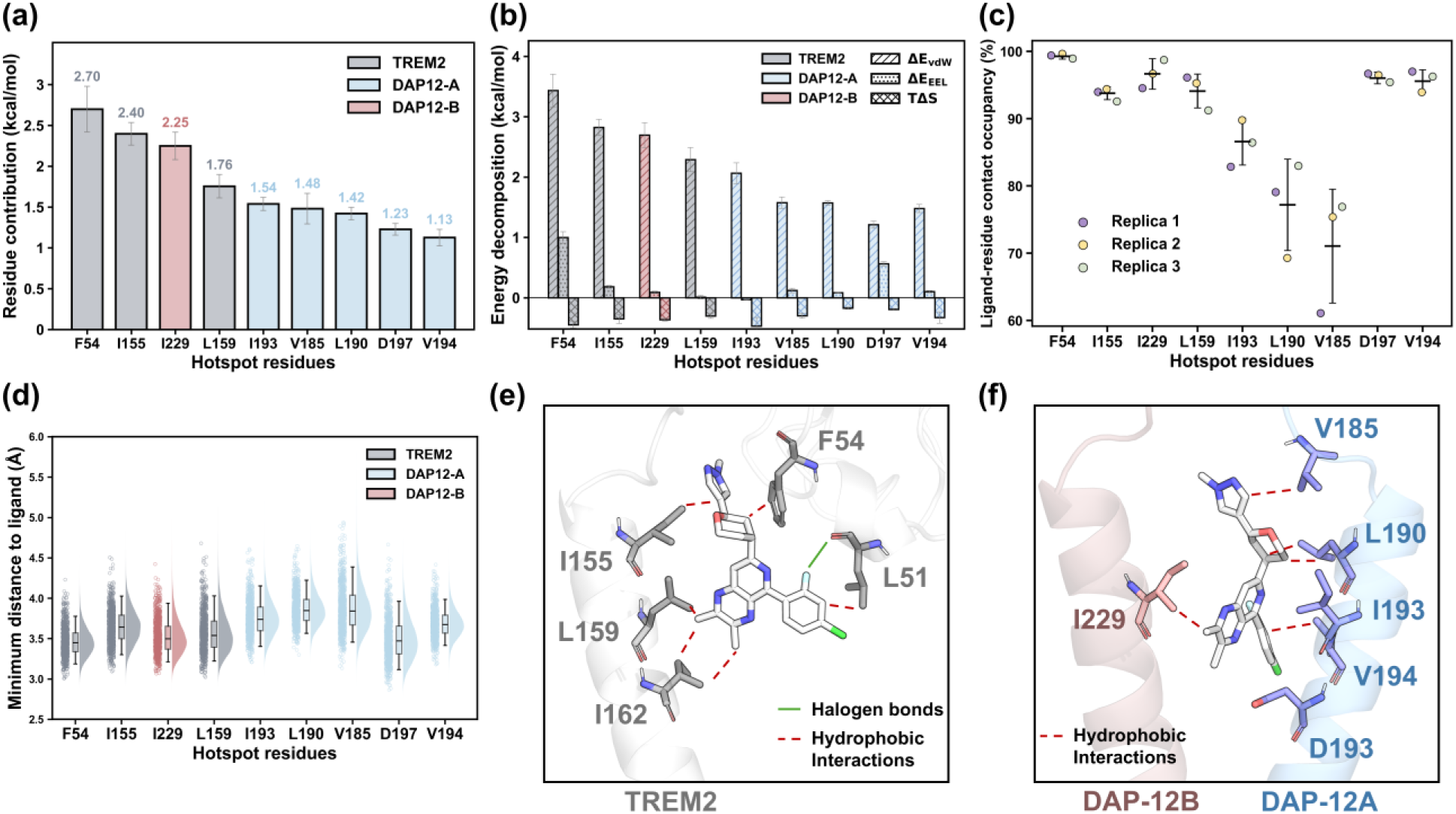
(a) Per-residue ASGBIE free-energy contributions of hotspot residues located within 5 Å of VG-3927. Only residues with contributions greater than 1 kcal·mol−1 are shown. Error bars represent the standard deviation across three independent replicas. Residues from TREM2, DAP12A, and DAP12B are colored gray, blue, and pink, respectively. (b) Energy decomposition of the nine hotspot residues contributing most strongly to VG-3927 binding, including van der Waals, electrostatic, polar solvation, and entropic terms. Different fill patterns indicate different energy components. (c) Ligand–residue contact occupancies of the hotspot residues across three independent replicas, shown in purple, yellow, and green. (d) Distributions of the minimum ligand–residue distances for the hotspot residues. Violin plots show the sampled distance distributions, with mean values and whiskers indicated. Residues from TREM2, DAP12A, and DAP12B are colored gray, blue, and pink, respectively. (e,f) Representative three-dimensional views of the hotspot interaction network surrounding VG-3927, showing TREM2-side interactions in panel (e) and DAP12-side interactions in panel (f).

Energy decomposition further showed that van der Waals interactions are the dominant energetic component of VG-3927 binding (**Figure 4b**). In contrast, electrostatic contributions were generally smaller and were partially offset by unfavorable polar solvation terms. This energetic profile is consistent with the nonpolar character of the TM cavity and the lipophilic nature of VG-3927. Notably, the contribution of D197 suggests that polar or charged residues may still assist ligand recognition through local packing, orientation effects, or transient polar contacts, even though the overall binding process is primarily driven by hydrophobic and dispersion interactions.

To further evaluate the persistence of these interactions, we analyzed ligand–residue contact occupancies and minimum ligand–residue distances (**Figure 4c,d**). A core anchoring layer composed of F54, I155, I229, and L159 maintained near-continuous contact with VG-3927, with occupancies generally above 90%. In contrast, I193, L190, V185, V194, and D197 formed a secondary contact network with slightly lower but still persistent occupancies, ranging from 70% to 95% in average. The corresponding minimum-distance distributions were narrowly centered around 3–4 Å, indicating that these residues remain in close proximity to the ligand throughout the simulations.

Representative protein–ligand interaction views further clarify the structural basis of VG-3927 recognition (**Figure 4e,f**). VG-3927 is surrounded mainly by hydrophobic side chains from TREM2 and DAP12, with additional polar contacts contributing to local orientation. Based on its chemical structure, VG-3927 can be divided into four functional fragments: a halogenated phenyl ring, a fused aza-aromatic core, an oxygen-containing six-membered ring, and a distal N-methyl aza-heterocycle.

The halogenated phenyl ring is accommodated along the DAP12-facing side of the cavity. It is surrounded by a hydrophobic wall formed by V185, L190, I193, and V194 from DAP12A, together with I229 from DAP12B. Although the halogen substituent may increase lipophilicity and participate in a halogen-bond-like interaction, the dominant stabilizing force in this region is likely close hydrophobic packing within the nonpolar pocket. The central fused aza-aromatic core functions as the main anchoring scaffold. Its rigid and planar structure provides an extended hydrophobic surface that is recognized by L51, F54, I155, I162, and L159 on TREM2. Among these residues, F54 is particularly important because its aromatic side chain can form favorable hydrophobic and aromatic packing interactions with the ligand scaffold, consistent with the highest free-energy contribution. The oxygen-containing six-membered ring appears to enhance three-dimensional shape complementarity by occupying space near I155, L159, and neighboring hydrophobic residues. Finally, the distal N-methyl aza-heterocycle likely acts as an auxiliary recognition group that helps define ligand orientation and stabilize the overall bound pose.

Taken together, these results indicate that VG-3927 is stabilized by a cooperative interaction network within the TREM2–DAP12 TM cavity. The hotspot residues are distributed across all three helices, allowing different regions of VG-3927 to participate in distinct interactions in the receptor–adaptor interface. This binding pattern is dominated by tight hydrophobic packing, supplemented by local polar contacts, and provides a structural explanation for how VG-3927 may stabilize the TREM2–DAP12 complex.

### 2.4 Effect of VG-3927 on TREM-DAP12 interaction

#### TREM-DAP12 residue contact map

To study the effect of VG-3927 on the TM protein–protein interface, we quantified residue-pair contact occupancies between the TREM2 helix and the two DAP12 helices in both ligand-free and ligand-bound simulations. For clarity, the ligand-free TREM2–DAP12 complex is referred to as the apo state, whereas the VG-3927-bound complex is referred to as the holo state. Contacts were defined using a 4 Å heavy-atom distance cutoff, and mean occupancies were averaged over three independent 200 ns replicas. Only contacts with occupancies above 50% in at least one state were retained, allowing the analysis to focus on reproducible structural features rather than transient fluctuations.

Comparison of the contact maps of apo and holo states (**Figure 5a,b**) shows that VG-3927 binding induces a selective reorganization of the interhelical contact network. In the apo state, the TREM2–DAP12A interface is more extensive than the TREM2–DAP12B interface, suggesting a preorganized asymmetric assembly. Upon VG-3927 binding, many pre-existing TREM2–DAP12A contacts are maintained and further reinforced, whereas the TREM2–DAP12B interface gains several new or strengthened contacts. On the DAP12A side, one major contact cluster is formed between the middle region of the TREM2 helix, particularly residues 163–166, and DAP12A residues 193–204. High-occupancy pairs such as 166–197, 166–201, and 166–204 define a central interaction core. A second cluster is observed between the upper region of TREM2 and DAP12A, involving contacts such as 173–207, 174–203, 174–207, and 176–211. On the DAP12B side, the holo state shows reinforced contacts including 162–230, 166–230, and 166–233. In addition, new ligand-associated contacts, including 154–226, 155–226, 158–226, and 166–237, appear in the holo simulations, while 158–230 is also strengthened. The emergence of the 159–193 contact between TREM2 and DAP12A further supports ligand-induced remodeling of the TM interface. These changes indicate that VG-3927 promotes a more integrated three-helix assembly by reinforcing the relatively weak DAP12B-side interface while preserving and strengthening the preorganized DAP12A-side contacts.

**Figure 5.**
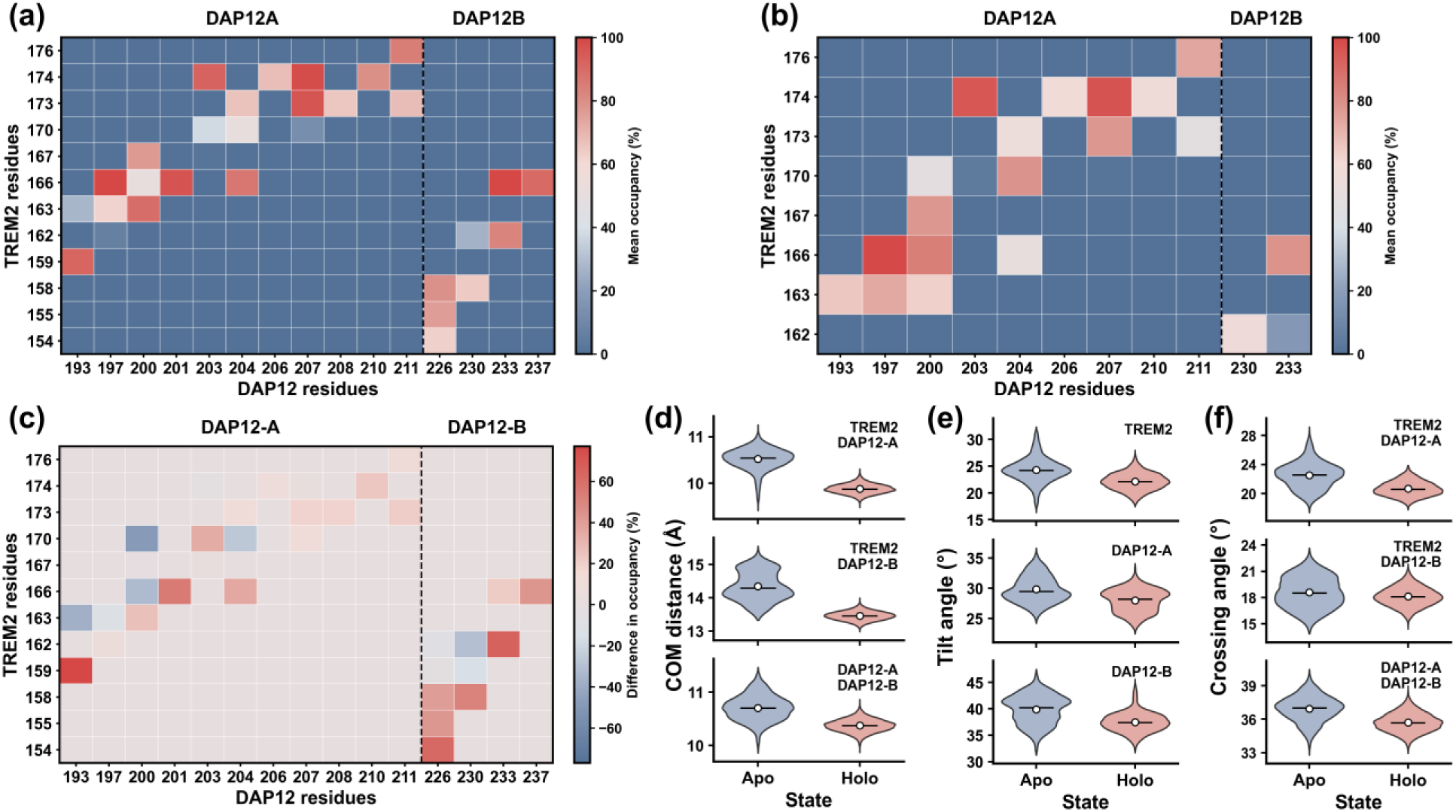
(a,b) Residue-pair contact occupancy maps of the TREM2–DAP12 TM interface in the apo and holo states, averaged over three independent replicas. TREM2 residues are shown on the y-axis, and DAP12A and DAP12B residues are shown on the x-axis. The black dashed line separates DAP12A and DAP12B residues. (c) Difference map of residue-pair contact occupancy between the holo and apo states, calculated as holo minus apo. Red indicates increased contact occupancy in the holo state, whereas blue indicates decreased contact occupancy. (d) Center-of-mass (COM) distance distributions for the three helix pairs in the apo and holo states. (e) Tilt-angle distributions of TREM2, DAP12A, and DAP12B in the apo and holo states. (f) Crossing-angle distributions for the three helix pairs in the apo and holo states.

The contact difference map (**Figure 5c**) further shows that ligand-induced remodeling is spatially asymmetric. Several apo-state contacts in the lower TREM2–DAP12A region decrease in occupancy, whereas multiple contacts in the upper TREM2–DAP12A region and on the TREM2–DAP12B side increase in the holo state. Thus, VG-3927 does not simply tighten the entire interface. Instead, it redistributes residue-level interactions across the three-helix assembly. This asymmetric remodeling is consistent with the hotspot analysis, in which DAP12A provides a broader ligand-contacting surface, whereas DAP12B contributes a more localized but important anchoring role. The gain of DAP12B-side contacts in the holo state suggests that asymmetric ligand binding is translated into broader interhelical stabilization, thereby coupling the bound ligand to the receptor–adaptor interface.

These residue-level changes are consistent with the helix-packing parameters shown in **Figure 5d–f**. In the holo state, the center-of-mass (COM) distances decrease for all three helix pairs compared with the apo state, indicating tighter interhelical packing upon ligand binding. The crossing angles also shift to smaller values, consistent with a more parallel arrangement of the helices. In addition, the tilt angles of TREM2, DAP12A, and DAP12B decrease in the holo state, indicating that the helices adopt a more upright orientation relative to the membrane normal. The holo-state distributions are also generally narrower than the corresponding apo-state distributions, suggesting reduced conformational heterogeneity and a more constrained TM assembly. Together, these geometric changes indicate that VG-3927 stabilizes a TM complex that is more compact, more ordered, and more geometrically coherent. Detailed information is provided in **Table S4**-**S5**.

We further quantified the occupancies of key salt-bridge like interactions and hydrogen bonds between the conserved TREM2 K26 residue and the two DAP12 helices. Residues involved in this network are described using the literature numbering to remain consistent with previous studies. TREM2 K26 is critical for receptor–adaptor assembly because it mediates electrostatic pairing with DAP12 acidic residues within the membrane. Across three independent replicas, the holo state consistently showed higher occupancies for these anchoring interactions than the apo state (**Table 1**).

**Table 1.**
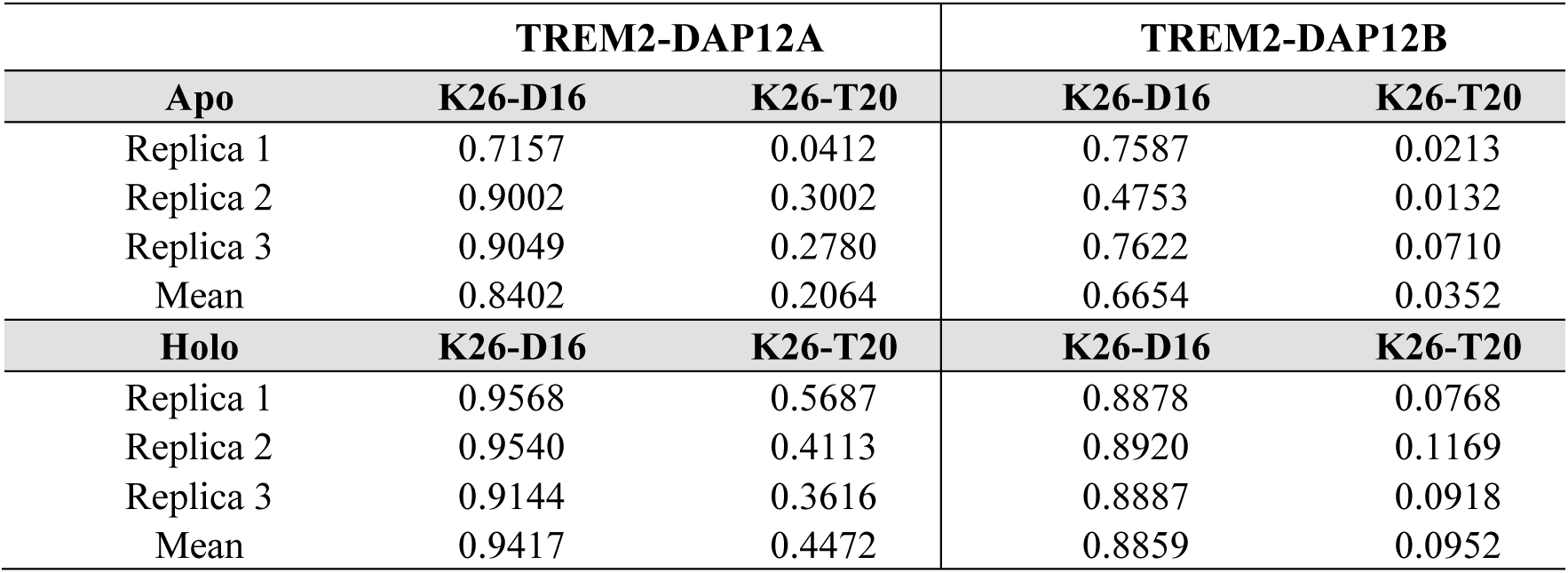
Hydrogen bonds occupancy of TM regions of TREM2 and DAP12 dimer.

At the TREM2–DAP12A interface, the mean occupancy of the K26–D16 interaction increased from 0.84 in the apo state to 0.94 in the holo state, whereas the K26–T20 interaction increased from 0.21 to 0.45. A similar trend was observed at the TREM2–DAP12B interface, where the K26–D16 occupancy increased from 0.67 to 0.89 and the K26–T20 occupancy increased from 0.04 to 0.10. These results indicate that VG-3927 promotes more persistent receptor–adaptor anchoring interactions at the membrane-embedded interface. In particular, the strengthening of the K26–D16 electrostatic contacts suggests that ligand binding stabilizes a more favorable assembly state, whereas the increased K26–T20 occupancies indicate improved local helix packing and side-chain alignment in the holo state.

Overall, VG-3927 binding within the TM cavity strengthens interhelical coupling and shifts the TREM2–DAP12 complex toward a more compact, upright, and ordered arrangement. The contact maps indicate a selective redistribution of residue-level interactions, while the geometric analyses and anchoring-network occupancies show that these local changes are associated with global stabilization of the TM assembly. Although these simulations do not directly measure downstream signaling, the observed reinforcement of the TREM2–DAP12 interface provides a structural basis for how VG-3927 may enhance receptor–adaptor coupling and promote signaling-competent assembly formation.

#### PCA analysis of TREM2–DAP12 complex

We next performed principal component analysis (PCA) on the apo and holo trajectories and compared their conformational sampling and free-energy landscapes (FES). PCA provides an ensemble-level description of the dominant collective motions of the three-helix bundle, allowing us to evaluate whether ligand binding mainly introduces a new conformational state or restricts the receptor–adaptor complex to a more defined subset of pre-existing conformations.

In the apo state, the three independent replicas occupied partially distinct regions within the PC1–PC2 subspace (**Figure 6a**). Replica 1 mainly sampled the upper-right region, replica 2 extended broadly toward negative PC1 values, and replica 3 explored a distinct region at positive PC1 and lower PC2 values. This incomplete overlap indicates that the ligand-free complex samples multiple conformational basins and retains substantial structural heterogeneity, suggesting that the two DAP12 helices can adopt different packing arrangements around the TREM2 helix.

**Figure 6.**
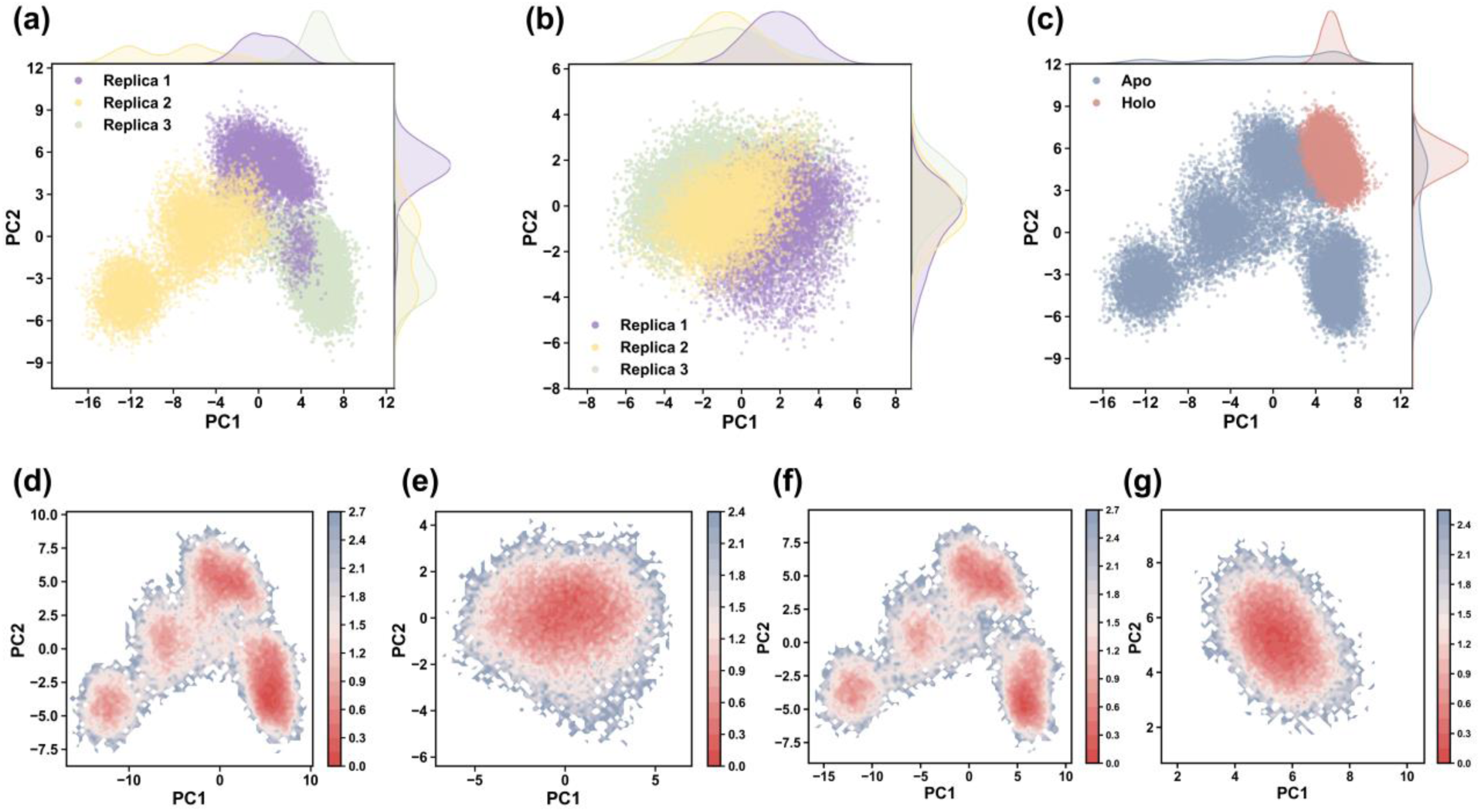
(a,b) Conformational sampling of the apo (a) and holo (b) states projected onto their respective first two principal components, PC1 and PC2. Three independent replicas are shown in purple, yellow, and green. (c) Comparative projection of the apo and holo ensembles onto the apo-derived PCA space, with the apo ensemble shown in blue and the holo ensemble shown in red. (d,e) Two-dimensional free-energy surfaces (FES) of the apo (d) and holo (e) states calculated from their respective PC1 and PC2 coordinates. (f,g) Free-energy landscapes of the apo (f) and holo (g) ensembles projected onto the common apo-derived PCA reference frame.

By contrast, the holo state showed much stronger spatial overlap among the three replicas and was concentrated within a more confined region of PCA space (**Figure 6b**). The marginal distributions along both PC1 and PC2 were also narrower than those in the apo state, indicating that VG-3927 reduces the amplitude of dominant collective motions. This convergence across independent simulations suggests that ligand binding facilitates replica-dependent convergence and stabilizes a more reproducible conformational ensemble. Therefore, VG-3927 does not merely remain bound within the TM cavity; it also constrains the global dynamics of the three-helix assembly.

To directly compare the two ensembles, the holo trajectories were projected onto the PCA space defined by the apo simulations (**Figure 6c**). In this common reference space, the holo ensemble occupied a restricted region within the broader apo conformational landscape. Importantly, the holo distribution did not extend into a completely separate region of PCA space, indicating that VG-3927 does not drive the complex into a wholly new conformational state. Instead, the ligand appears to select and stabilize a pre-existing conformational substate that is already accessible to the apo complex.

The FES analysis further supports this interpretation. The apo FES displayed multiple local minima distributed across PC1 and PC2 (**Figure 6d,f**), indicating that the ligand-free complex can convert between several metastable conformational states. These minima likely reflect alternative helix-packing arrangements within the TREM2–DAP12 helices. In contrast, the holo FES was dominated by a single, narrower low-energy basin (**Figure 6e,g**). The reduction in the number and spatial distribution of low-energy basins indicates that VG-3927 decreases conformational heterogeneity and reshapes the energy landscape toward a more constrained state. This landscape narrowing is consistent with reduced configurational entropy, which is expected when a ligand stabilizes a compact receptor–adaptor assembly.

These PCA and FES results provide an ensemble-level explanation for the residue-contact and geometric changes described above. The strengthened interhelical contacts, reduced COM distances, smaller crossing angles, and more upright helix orientations are not isolated local features. Rather, they reflect a ligand-induced redistribution of the conformational landscape toward a compact and ordered substate. By bridging the TREM2 and DAP12 helices within the TM cavity, VG-3927 restricts alternative helix-packing modes and stabilizes a coherent three-helix bundle. This conformational focusing provides a plausible structural basis for enhanced receptor–adaptor coupling and supports the molecular-glue-like mechanism proposed for VG-3927. Overall, VG-3927 binding appears to act at two coupled levels: locally, by engaging hotspot residues across the TREM2–DAP12 interface, and globally, by restricting the conformational ensemble to a more stable and signaling-compatible transmembrane architecture.

#### DCCM analysis of TREM2–DAP12 complex

Dynamic cross-correlation matrix (DCCM) analysis was performed to further reveal how VG-3927 affects the internal dynamics of the TREM2–DAP12 complex. DCCM describes the correlated and anticorrelated motions between residue pairs and therefore provides information complementary to contact maps and geometric analyses. Here, both residue-level and domain-averaged correlation patterns were compared between the apo and holo states.

In the apo state, correlated motions were mainly localized within individual structural regions, particularly within the TREM2 Ig-like domain and within parts of the membrane-embedded helices (**Figure 7a**). However, the coupling between different regions was relatively heterogeneous, suggesting that the ligand-free complex retains substantial dynamic flexibility. This pattern is consistent with the PCA and contact-map analyses, which showed that the apo complex samples multiple conformational substates and contains a less integrated TREM2–DAP12B interface.

**Figure 7.**
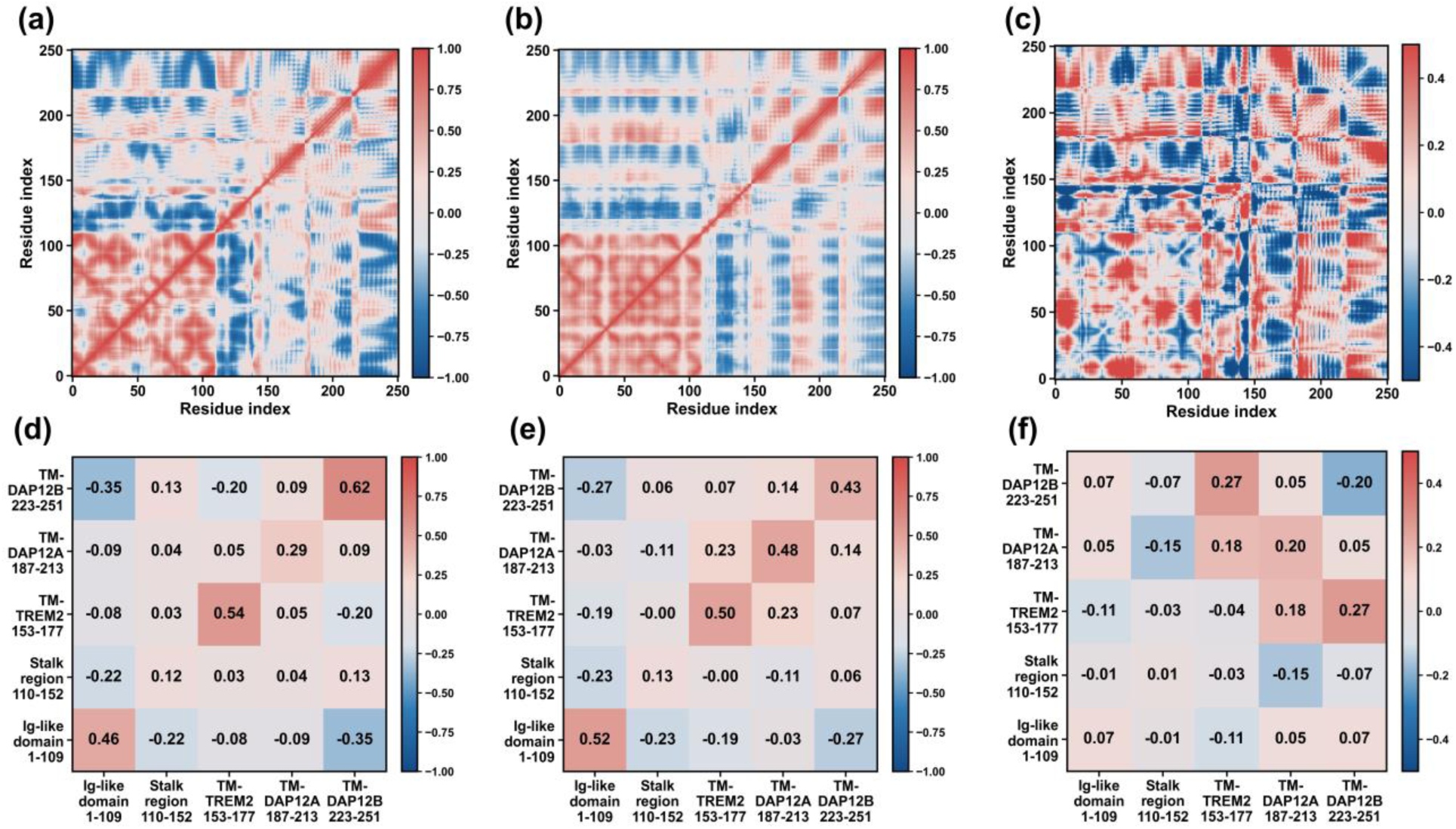
(a, b) Residue-resolved dynamic cross-correlation matrices (DCCM) for the apo (a) and holo (b) states, illustrating the internal signaling dynamics of the receptor–adaptor assembly. (c) Difference DCCM map calculated as DCCM_holo_ - DCCM_apo_; red regions indicate enhanced cooperative correlation, while blue regions indicate diminished correlation in the holo state; (d, e) Domain-averaged correlation matrices for the apo (d) and holo (e) states, partitioned into five functional regions: TREM2 Ig-like domain (residues 1–109), stalk region (residues 110–152), TREM2 transmembrane (TM) helix (residues 153–177), DAP12-A TM helix (residues 187–213), and DAP12-B TM helix (residues 223–251). (f) Difference map of the domain-averaged correlations, highlighting the selective enhancement of cooperative motion within the transmembrane core upon VG-3927 binding.

In contrast, the holo state displayed a more organized correlation pattern within the TM core and a spatially asymmetric trend upon ligand binding (**Figure 7b**). Increased positive correlations are mainly observed at the interfaces connecting the TREM2 TM helix with DAP12A and DAP12B. The most obvious changes involve the central and upper regions of the TM core, spanning TREM2 residues 160–176, DAP12A residues 195–210, and DAP12B residues 228–240. These regions overlap with the sites that showed strengthened contact occupancies and holo-specific interface remodeling in the previous analyses. Thus, VG-3927 not only stabilizes local residue contacts but also increases the dynamic coupling of these interhelical regions.

Domain-averaged DCCM analysis provides a clearer view of this effect (**Figure 7d–f**). The largest ligand-induced increase in positive correlation occurs between the TREM2 TM helix, residues 153–177, and DAP12B, residues 223–251, with the averaged cross-correlation coefficient increasing by approximately +0.27. A smaller but still clear increase is observed between the TREM2 TM helix and DAP12A, with an enhancement of approximately +0.18. By contrast, the correlation between the two DAP12 helices increases only modestly, by approximately +0.05. These results indicate that VG-3927 enhances cooperative motion between TREM2 and each DAP12 helix, with the strongest effect on the DAP12B side.

This finding is particularly informative when compared with the residue-level binding analysis. The direct energetic hotspots of VG-3927 are concentrated mainly on TREM2 and DAP12A, whereas DAP12B contributes fewer direct ligand-contacting residues. However, DAP12B displays the largest increase in dynamic correlation with TREM2 upon ligand binding. This apparent discrepancy may arise because VG-3927 first stabilizes the TREM2–DAP12A interface and subsequently propagates this stabilizing effect to the previously weaker TREM2–DAP12B interface. In this model, DAP12B acts as a major recipient of ligand-induced dynamic stabilization, even though it does not constitute the dominant direct binding surface.

Ligand binding also redistributes longer-range dynamic coupling within the receptor–adaptor complex. The correlation between the TREM2 TM helix and the Ig-like domain decreases by approximately -0.11, whereas the coupling between the TM helix and the stalk region remains largely unchanged. These changes suggest that VG-3927 selectively strengthens coordinated motions within the TM core while partially reducing the influence of distal extracellular fluctuations on the TM region. Such selective dynamic focusing is consistent with the PCA results, in which the holo ensemble occupied a narrower conformational basin without forming a completely new state.

Taken together, the DCCM analysis supports a model in which VG-3927 converts local transmembrane binding into broader dynamic stabilization of the TREM2–DAP12 assembly. The ligand enhances cooperative motion specifically within the membrane-embedded receptor–adaptor core, with the largest dynamic gain occurring between TREM2 residues 153–177 and DAP12B residues 223–251 and a parallel enhancement between TREM2 residues 153–177 and DAP12A residues 187–213. These dynamic changes complement the strengthened contact occupancies, reduced helix–helix distances, and narrowed PCA/FES landscapes described above. Therefore, VG-3927 appears to stabilize the complex not only by forming direct hydrophobic contacts in the TM cavity but also by promoting a more coordinated and dynamically coupled three-helix architecture. This provides a mechanistic explanation for how a locally bound lipophilic ligand may reinforce receptor–adaptor coupling at the membrane interface.

## 3 SUMMARY AND DISCUSSION

In this study, we constructed a structural and dynamic model to explain how the clinical-stage small-molecule TREM2 agonist VG-3927 may stabilize the membrane-associated TREM2–DAP12 complex. By integrating AlphaFold 3-based structural modeling, blind docking, multi-replica all-atom MD simulations, residue-level free-energy decomposition, interhelical contact analysis, helix-packing measurements, PCA/FES analysis, and DCCM analysis, we identified a plausible membrane-embedded binding mode in which VG-3927 occupies an interfacial cavity formed by the transmembrane helices of TREM2, DAP12 dimer.

The simulations suggest that this cavity-bound pose is stable and reproducible across independent replicas. VG-3927 binding is supported by a focused hotspot network dominated by TREM2 and DAP12A residues, with an additional contribution from DAP12B. The interaction profile is mainly governed by van der Waals and hydrophobic packing interactions, consistent with the lipophilic character of VG-3927 and the nonpolar environment of the transmembrane cavity.

Beyond direct protein–ligand recognition, VG-3927 binding is associated with broader remodeling of the TREM2–DAP12 transmembrane assembly. The holo complex retains the major apo-state scaffold but shows selective reorganization of residue-level contacts, including reinforcement of TREM2–DAP12A interactions and the formation or strengthening of contacts on the TREM2–DAP12B side. These local changes are accompanied by reduced interhelical center-of-mass distances, smaller crossing angles, lower tilt angles, and increased occupancies of conserved TREM2 K26–DAP12 D16/T20 anchoring interactions. Together, these features indicate that ligand binding shifts the three-helix assembly toward a more compact, upright, and ordered state.

Ensemble-level analyses further support this mechanism. PCA and free-energy landscapes show that the apo complex samples multiple conformational basins, whereas the VG-3927-bound complex occupies a narrower and more reproducible conformational subspace. Projection of the holo ensemble onto the apo-defined PCA space suggests that VG-3927 does not create a completely new global state, but instead selects and stabilizes a pre-existing transmembrane substate. DCCM analysis indicates that this stabilization is accompanied by enhanced cooperative motion within the membrane-embedded core, especially between the TREM2 transmembrane helix and DAP12B. Thus, local ligand binding appears to propagate into global dynamic stabilization of the receptor–adaptor interface.

Because the proposed binding site and residue contributions are derived solely from computational modeling, they should be considered testable mechanistic hypotheses rather than established facts. Future experimental validation—through mutagenesis, ligand structure–activity relationship studies, and structural approaches such as cryo-EM or X-ray crystallography—will be essential to confirm the predicted hotspot residues and to assess whether transmembrane interfacial stabilization represents a general strategy for developing next-generation small-molecule TREM2 agonists.

In summary, the computational predictions presented in this study offer a valuable starting point for understanding the transmembrane mode of action of VG-3927. our results support a model in which VG-3927 acts as a membrane-adapted interfacial stabilizer of the TREM2–DAP12 complex. By occupying a three-helix transmembrane cavity, the ligand reinforces local hotspot interactions, improves receptor–adaptor helix packing, and promotes a more dynamically coupled assembly that is compatible with enhanced signaling. However, given the limitations discussed above, the model must be complemented by experimental validation before it can serve as a structural pharmacological foundation for TREM2 agonist activity.

## 4 METHODOLOGY

### Apo Structure Preparation of the TREM2–DAP12 Complex

The amino acid sequences of human TREM2 and DAP12 were retrieved from UniProt,^24^ and the receptor–adaptor complex was predicted using AlphaFold 3^25^. The predicted full-length assembly was trimmed before molecular dynamics (MD) simulations to reduce computational cost and to focus on the membrane-associated receptor–adaptor interface.

The resulting apo TREM2–DAP12 model was oriented relative to the membrane using the PPM 2.0 module implemented in the OPM server,^26^ which estimates membrane placement based on hydrophobic thickness and transfer energetics. The membrane-oriented system was then constructed using PACKMOL-Memgen^27^. A mixed bilayer composed of POPC and cholesterol at a molar ratio of 80:20 was used to approximate a cholesterol-containing eukaryotic membrane environment. The system was solvated with TIP3P water, and Na+ and Cl− ions were added to neutralize the system and reach a salt concentration of 0.15 M. A water layer of 20.0 Å was added on each side of the membrane to ensure sufficient hydration of the protein–membrane complex.

The AMBER ff14SB force field was used for the protein^28^, Lipid17 was used for POPC and cholesterol^29^, and the TIP3P^30^ model was used for water molecules. The apo TREM2–DAP12 complex was simulated for 200 ns. After the simulation, clustering analysis was performed to identify a representative and structurally stable receptor conformation, which was subsequently used for ligand docking.

### Holo Structure Preparation of the VG-3927/TREM2–DAP12 Complex

VG-3927 was obtained in SDF format from PubChem^31^ and geometry-optimized using Gaussian 16^32^. The optimized ligand structure was then used for docking and MD system preparation. Ligand force-field parameters were generated using GAFF2^33^, and atomic partial charges were assigned using RESP.

Blind docking was performed using AutoDock Vina^34^ with the representative apo-state receptor conformation as the docking receptor. The docking grid box was defined to cover the full membrane-associated receptor assembly, including the transmembrane interface, so that no binding site was imposed a priori. Among the cavity-bound poses, a top-ranked pose with a docking score of −9.6 kcal·mol^−1^ was selected for further evaluation. This pose was not chosen solely according to the docking score; visual inspection and a short preliminary MD assessment were also used to evaluate whether the ligand remained stably located within the transmembrane cavity. The selected pose was then used as the initial model for holo-state membrane simulations.

The holo system was constructed using the same membrane-building procedure, lipid composition, solvation protocol, ion concentration, and force-field settings as those used for the apo system.

### Molecular Dynamics Simulations

All apo and holo systems were simulated using the same general MD workflow in three independent replicas. The starting coordinates were taken from the corresponding membrane-embedded, solvated, and ionized systems containing the explicit lipid bilayer, water molecules, and ions. The pmemd module of AMBER22 was employed.^35^

Energy minimization was carried out in three stages. First, all atoms except water molecules and ions were restrained with a harmonic force constant of 100 kcal·mol^−1^·Å^−2^, and the system was minimized for 50,000 steps to relax solvent and lipid packing around the protein complex. Second, the restraint force constant was reduced to 2 kcal·mol^−1^·Å^−2^, followed by 20,000 steps of minimization. Third, all restraints were removed, and an additional 20,000 steps of unrestrained minimization were performed to relax the full system.

After minimization, each system was gradually heated from 0 to 310 K over 1 ns. This was followed by a three-stage equilibration protocol in which positional restraints on the solute were progressively reduced from 5 to 2 to 0 kcal·mol^−1^·Å^−2^. Each equilibration stage was run for 2 ns, allowing the protein–membrane system to adapt gradually to production conditions. Production simulations were then performed for 200 ns in the NPT ensemble. The first 20 ns of each production trajectory were discarded as equilibration before analysis.

A nonbonded cutoff of 10 Å was used for all minimization, heating, equilibration, and production simulations. Long-range electrostatic interactions were treated using the particle mesh Ewald (PME) method.^36^ Bonds involving hydrogen atoms were constrained using the SHAKE algorithm.^37^ Because the systems contained an explicit membrane bilayer, anisotropic pressure coupling with surface-tension correction was applied using csurften = 3 to maintain appropriate bilayer behavior during production simulations.

For the holo system, an additional relaxation step was introduced before initiating the three independent production replicas. After construction of the ligand-bound complex, a short 10 ns production simulation was performed, and a representative stable structure from this trajectory was extracted as the common starting conformation for the three independent replicas. This procedure allowed the ligand-bound system to relax locally before formal production simulations.

After MD simulations, conformational clustering was performed on the holo trajectories to identify representative ligand-bound conformations. Conformational clustering was performed using CPPTRAJ to identify representative VG-3927-bound poses from the holo trajectories. Clustering was based on the heavy atoms of VG-3927, excluding hydrogen atoms, and was carried out using a hierarchical agglomerative algorithm with 10 clusters and an RMSD cutoff of 2.0 Å. To reduce computational cost, every 10th frame was used for clustering with random sieving. The representative structure from the most populated cluster of each replica was selected as the dominant ligand-bound conformation for that trajectory. In addition, the three holo trajectories were concatenated and subjected to the same clustering protocol to obtain an overall representative VG-3927-bound conformation.

Post-simulation analyses, including RMSD, RMSF, residue-contact occupancy, minimum ligand–residue distances, helix center-of-mass distances, crossing angles, and tilt angles, were performed using CPPTRAJ in AMBER 22. Contacts were defined using a 4 Å heavy-atom distance cutoff unless otherwise stated.

### Principal Component Analysis and Free-Energy Surface Construction

Principal component analysis (PCA) was performed using CPPTRAJ in AMBER 22 to characterize the dominant collective motions of the apo and holo TREM2–DAP12 TM assemblies. For each system, three trajectories were combined and aligned to a reference structure using Cα atoms to remove overall translational and rotational motions. To avoid the influence of early equilibration, frames from 20 to 200 ns of each trajectory were used for PCA calculation.

For the separate apo and holo PCA analyses, each state was projected onto its own eigenvector space derived from the corresponding covariance matrix. To directly compare conformational sampling between the two states, the holo trajectories were also projected onto the apo-derived PCA space. Two-dimensional free-energy surfaces were constructed from the probability distribution of the projected PC1 and PC2 coordinates. The minimum free energy was set to zero for visualization.

### Dynamic Cross-Correlation Matrix Analysis

Dynamic cross-correlation matrix (DCCM) analysis was performed using CPPTRAJ in AMBER 22 to evaluate correlated and anticorrelated residue motions in the apo and holo TREM2–DAP12 complexes. For each state, the three independent production trajectories were combined for analysis. To avoid the influence of early equilibration, frames from 20 to 200 ns of each trajectory were used for DCCM calculation. DCCM was calculated using the Cα atoms of residues 1–251.

For domain-level analysis, residue-wise correlation coefficients were averaged over predefined structural regions, including the Ig-like domain, stalk region, TREM2 transmembrane helix, DAP12A transmembrane helix, and DAP12B transmembrane helix.

### ASGBIE calculation of free energies

Residue-level binding free-energy decomposition was performed using alanine scanning with generalized Born and interaction entropy (ASGBIE). This method combines MM/GBSA based on the generalized Born (GB) solvent model with interaction entropy (IE) to estimate the energetic contribution of individual residues by computationally mutating each selected residue to alanine. The total residue contribution, including both enthalpic and entropic components, was calculated as the binding free-energy difference between the wild-type residue (w) and its alanine-mutated counterpart (a), as defined below,^38–40^

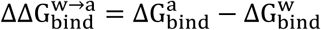

where 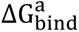 and 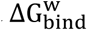 are the binding free energies of the alanine-mutant and the wild-type complex, respectively. The binding free energy is decomposed into several components:

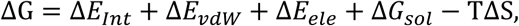

where Δ*E_Int_*, Δ*E_vdW_*, Δ*E_ele_*, Δ*E_sol_*, and TΔS represent, the internal energy, the van der Waals interaction, the electrostatic interaction, the solvation free energy, and the entropy contributions. The solvation free energy was further expressed as the sum of polar and nonpolar solvation terms:

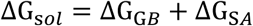

where ΔG_G*B*_is the polar solvation free energy calculated using the generalized Born model, and ΔG_S*A*_is the nonpolar solvation free energy estimated from the solvent-accessible surface area (SASA), as defined below,

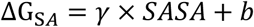

where *γ* and *b* are empirical parameters.

Therefore, the total binding free energy at the binding site can be expressed as the sum of the individual residue contributions:^41,42^

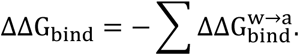

The entropic component, which is often difficult to estimate accurately in conventional MM/GBSA calculations, was evaluated using the interaction entropy method, as defined below,^42^

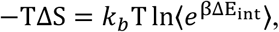

where β = 1/*k_b_T*, and ΔE_int_ denotes the fluctuation in the receptor–ligand interaction energy, and 〈… 〉 is the ensemble average over the MD trajectory.

The ASGBIE analysis was performed based on the initial 200 ns MD trajectories of the VG-3927-bound TREM2–DAP12 complex. To ensure the system had reached structural equilibrium, a stable segment spanning 100–120 ns was extracted for the free-energy calculations. These analyses utilized the Generalized Born (GB) model with the igb=2 setting, incorporating a physiological salt concentration of 0.15 M to accurately simulate the ionic environment. To identify the critical hotspots stabilizing the ligand, individual energetic contributions were quantified for all amino acid residues situated within a 5 Å radius of the binding pocket.

## SUPPORTING INFORMATION

The supporting information includes the binding free energy of each hotspot residue of the VG-3927 and TREM-DAP12 complex; energy decomposition and standard deviation of each hotspot residue; ligand–residue contact occupancies of the main hotspot residues; tilt angle, crossing angle and center-of-mass (COM) distance of TREM2-DAP12 apo complex and VG-3927/TREM2-DAP12 holo complex.

## ACKNOWLEDGEMENT

This work was supported by the National Natural Science Foundation of China (Grant nos. 22333006, 92270001), Shanghai Municipal Science and Technology Commission (Grant No. 25DX2800500), Guangdong Provincial Pearl River Talent Program (Grant No. 2023ZT10L045). We also acknowledge the support by NYU High Performance Computing resources and ECNU Multifunctional Platform for Innovation (001).

