## Supplemental Data 1 for "Structural and Dynamic Basis of TREM2–DAP12 Stabilization by Small-Molecule Agonist VG-3927"

**Table S1**. Hotspot residues of the VG-3927/TREM2–DAP12 complex. dVDW, dEEL, dGB, dH, dIE and dG are the change in van der Waals contribution, electrostatic contribution, polar contribution, entropy, enthalpy, and binding free energy, respectively. The results are averaged over three independent simulations.

| Mut-Wid | dVDW | dEEL | dGB | dH | dIE | dG |
|---|---|---|---|---|---|---|
| **54PHE** | 3.4338 | 0.9985 | -1.5261 | 3.1465 | -0.4475 | 2.6990 |
| **155ILE** | 2.8228 | 0.1819 | -0.4791 | 2.7468 | -0.3503 | 2.3965 |
| **229ILE** | 2.6955 | 0.0903 | -0.4243 | 2.6122 | -0.3628 | 2.2494 |
| **159LEU** | 2.2898 | 0.0098 | -0.4236 | 2.0582 | -0.3028 | 1.7554 |
| **193ILE** | 2.0651 | -0.0320 | -0.1601 | 2.0079 | -0.4699 | 1.5380 |
| **185VAL** | 1.5763 | 0.1206 | -0.0253 | 1.7783 | -0.2974 | 1.4808 |
| **190LEU** | 1.5723 | 0.0836 | -0.1665 | 1.5935 | -0.1726 | 1.4209 |
| **197ASP** | 1.2118 | 0.5624 | -0.4984 | 1.4207 | -0.1931 | 1.2276 |
| **194VAL** | 1.4792 | 0.1007 | -0.2681 | 1.4537 | -0.3276 | 1.1261 |
| **198LEU** | 0.9376 | 0.0867 | -0.1134 | 1.0051 | -0.1175 | 0.8876 |
| **233ASP** | 0.6571 | 0.4943 | -0.4155 | 0.7801 | -0.0675 | 0.7126 |
| **151PRO** | 0.6459 | -0.0157 | 0.0421 | 0.6959 | -0.0968 | 0.5991 |
| **158LEU** | 0.7579 | 0.0041 | -0.1163 | 0.6903 | -0.1103 | 0.5800 |
| **51LEU** | 0.7514 | -0.0054 | -0.1408 | 0.6892 | -0.1299 | 0.5593 |
| **152PRO** | 0.7768 | 0.0359 | -0.1347 | 0.7252 | -0.1849 | 0.5403 |
| **162ILE** | 0.7612 | -0.1710 | 0.0472 | 0.7059 | -0.1803 | 0.5257 |
| **221VAL** | 0.5890 | 0.0157 | -0.1460 | 0.5190 | -0.0952 | 0.4238 |
| **163PHE** | 0.4896 | -0.0781 | 0.0868 | 0.5118 | -0.0881 | 0.4237 |
| **67ASP** | 0.1946 | 0.2711 | -0.2657 | 0.2332 | -0.0372 | 0.1960 |
| **230VAL** | 0.1284 | -0.0086 | 0.0087 | 0.1316 | -0.0076 | 0.1240 |
| **226LEU** | 0.1266 | 0.0285 | -0.0411 | 0.1140 | -0.0032 | 0.1109 |
| **184THR** | 0.0655 | 0.0664 | -0.0507 | 0.0814 | -0.0046 | 0.0768 |
| **65THR** | 0.0538 | -0.0190 | 0.0327 | 0.0675 | -0.0004 | 0.0671 |
| **66ASP** | 0.0122 | 0.0655 | -0.0509 | 0.0268 | -0.0016 | 0.0252 |
| **195MET** | 0.0249 | -0.0061 | -0.0013 | 0.0174 | -0.0011 | 0.0163 |
| **220THR** | 0.0087 | -0.0196 | 0.0150 | 0.0041 | -0.0002 | 0.0039 |
| **219SER** | 0.0012 | 0.0048 | -0.0074 | -0.0016 | -0.0001 | -0.0017 |
| **TOTAL** | 26.1291 | 2.8652 | -5.2230 | 25.8148 | -4.0504 | 21.7644 |

**Table S2**. Standard deviation (std) of hotspot residues of the VG-3927/TREM2–DAP12 complex. The results are averaged over three independent simulations.

| Mut-Wid | std_dVDW | std_dEEL | std_dGB | std_dH | std_dIE | std_dG |
|---|---|---|---|---|---|---|
| **54PHE** | 0.2708 | 0.0956 | 0.0421 | 0.2327 | 0.0546 | 0.2802 |
| **155ILE** | 0.1343 | 0.0221 | 0.0433 | 0.0998 | 0.0772 | 0.1386 |
| **229ILE** | 0.2064 | 0.0175 | 0.0410 | 0.1817 | 0.0265 | 0.1678 |
| **159LEU** | 0.1999 | 0.0286 | 0.0539 | 0.1401 | 0.0389 | 0.1427 |
| **193ILE** | 0.1713 | 0.0071 | 0.0265 | 0.1616 | 0.1446 | 0.0809 |
| **185VAL** | 0.0948 | 0.0368 | 0.0251 | 0.1493 | 0.0417 | 0.1872 |
| **190LEU** | 0.0360 | 0.0093 | 0.0210 | 0.0677 | 0.0176 | 0.0763 |
| **197ASP** | 0.0581 | 0.0369 | 0.0235 | 0.0703 | 0.0156 | 0.0723 |
| **194VAL** | 0.0674 | 0.0179 | 0.0115 | 0.0536 | 0.0944 | 0.1010 |
| **198LEU** | 0.0870 | 0.0031 | 0.0144 | 0.0843 | 0.0111 | 0.0803 |
| **233ASP** | 0.0426 | 0.0161 | 0.0115 | 0.0407 | 0.0245 | 0.0588 |
| **151PRO** | 0.1569 | 0.0120 | 0.0240 | 0.1702 | 0.0247 | 0.1671 |
| **158LEU** | 0.0361 | 0.0074 | 0.0203 | 0.0258 | 0.0346 | 0.0591 |
| **51LEU** | 0.3066 | 0.0075 | 0.0739 | 0.2870 | 0.0487 | 0.2465 |
| **152PRO** | 0.2499 | 0.0173 | 0.0240 | 0.2236 | 0.0388 | 0.2001 |
| **162ILE** | 0.0331 | 0.0035 | 0.0069 | 0.0358 | 0.0502 | 0.0858 |
| **221VAL** | 0.2501 | 0.0165 | 0.0917 | 0.2022 | 0.0449 | 0.1600 |
| **163PHE** | 0.1958 | 0.0436 | 0.0696 | 0.2280 | 0.0269 | 0.2517 |
| **67ASP** | 0.0727 | 0.0422 | 0.0429 | 0.0830 | 0.0272 | 0.0557 |
| **230VAL** | 0.0349 | 0.0051 | 0.0047 | 0.0342 | 0.0045 | 0.0298 |
| **226LEU** | 0.0203 | 0.0039 | 0.0095 | 0.0149 | 0.0009 | 0.0142 |
| **184THR** | 0.0066 | 0.0128 | 0.0136 | 0.0073 | 0.0017 | 0.0077 |
| **65THR** | 0.0048 | 0.0048 | 0.0073 | 0.0074 | 0.0001 | 0.0074 |
| **66ASP** | 0.0007 | 0.0113 | 0.0080 | 0.0040 | 0.0002 | 0.0042 |
| **195MET** | 0.0029 | 0.0156 | 0.0152 | 0.0033 | 0.0002 | 0.0031 |
| **220THR** | 0.0033 | 0.0125 | 0.0146 | 0.0056 | 0.0000 | 0.0057 |
| **219SER** | 0.0001 | 0.0047 | 0.0045 | 0.0003 | 0.0000 | 0.0003 |
| **TOTAL** | 0.7931 | 0.1409 | 0.3409 | 0.6512 | 0.0666 | 0.5846 |

**Table S3**. Ligand–residue contact occupancies of the hotspot residues across three independent replicas.

| Residue | Replica1 | Replica2 | Replica3 | Average |
|---|---|---|---|---|
| **F54** | 99.32 | 99.64 | 98.9 | 99.28 |
| **D197** | 96.97 | 97.22 | 98.13 | 97.44 |
| **V194** | 96.88 | 96.45 | 95.37 | 96.23 |
| **L159** | 96.16 | 95.59 | 94.94 | 95.56 |
| **I229** | 94.69 | 94.28 | 92.34 | 93.77 |
| **I155** | 94.26 | 94.07 | 91.92 | 93.41 |
| **I193** | 83.68 | 89.36 | 84.78 | 85.94 |
| **L190** | 79.54 | 74.07 | 81.5 | 78.37 |
| **V185** | 61.97 | 71.12 | 72.46 | 68.51 |

**Table S4**. Tilt angle, crossing angle and center-of-mass (COM) distance of TREM2-DAP12 apo complex. In this context, A, B and C represent TREM2, DAP12-A and DAP12-B.

| | Replica1 | Replica2 | Replica3 | Average |
|---|---|---|---|---|
| **A_tilt** | 26.8741 | 20.2024 | 26.3719 | 24.4828 |
| **B_tilt** | 35.6380 | 26.4339 | 27.8406 | 29.9708 |
| **C_tilt** | 40.3329 | 36.4999 | 43.3566 | 40.0631 |
| **AB_crossing** | 21.8322 | 20.3155 | 24.5147 | 22.2208 |
| **AC_crossing** | 19.1562 | 20.1036 | 17.2964 | 18.8521 |
| **BC_crossing** | 38.5109 | 38.0150 | 34.7027 | 37.0762 |
| **AB_COM_distance** | 10.5503 | 10.6792 | 9.8437 | 10.3577 |
| **AC_COM_distance** | 14.0462 | 14.6838 | 13.4270 | 14.0523 |
| **BC_COM_distance** | 10.7267 | 10.9629 | 11.1240 | 10.9379 |

**Table S5**. Tilt angle, crossing angle and center-of-mass (COM) distance of VG-3927/TREM2-DAP12 holo complex. In this context, A, B and C represent TREM2, DAP12-A and DAP12-B.

| | Replica1 | Replica2 | Replica3 | Average |
|---|---|---|---|---|
| **A_tilt** | 26.9602 | 19.7302 | 19.7009 | 22.1304 |
| **B_tilt** | 31.4443 | 30.0065 | 22.5278 | 27.9929 |
| **C_tilt** | 43.3188 | 33.5039 | 35.6411 | 37.4879 |
| **AB_crossing** | 19.6298 | 21.0504 | 21.3564 | 20.6789 |
| **AC_crossing** | 19.5848 | 17.9770 | 16.8148 | 18.1255 |
| **BC_crossing** | 36.3251 | 35.8852 | 34.9143 | 35.7082 |
| **AB_COM_distance** | 9.9160 | 9.8157 | 9.0967 | 9.6095 |
| **AC_COM_distance** | 13.2430 | 13.3662 | 13.8905 | 13.4999 |
| **BC_COM_distance** | 10.4516 | 10.7069 | 10.3736 | 10.5107 |
